# Perceptual learning reformats working memory codes along a reverse visual hierarchy

**DOI:** 10.64898/2026.03.20.712287

**Authors:** Siyuan Cheng, Yiran Ge, Nihong Chen

## Abstract

Mastering fine perceptual skills requires maintaining precise visual information while simultaneously processing new inputs. In the visual cortex, sensory and mnemonic representations coexist through region-wide multiplexing, yet how perceptual experience reshapes these neural codes remains unknown. Here, combining five days of intensive perceptual training with fMRI and multivariate pattern analyses, we tracked how perceptual learning reformats mnemonic neural representations in a motion-direction discrimination task. Learning improved motion-direction discrimination sensitivity and reshaped the distribution of sensory- and mnemonic-format codes across the visual hierarchy. Sensory-format maintenance was enhanced in V1, and the magnitude of this enhancement closely tracked the behavioral learning effect across individuals, whereas mnemonic-format coding was reduced in V3A. Within V1, learning further increased temporal coupling between sensory and mnemonic codes while leaving their near-orthogonal coding geometry largely unchanged. Our results suggest that perceptual learning reformats working memory codes along a reverse visual hierarchy, strengthening high-fidelity sensory-like representations while preserving the separable organization of multiplexed codes. These findings identify reverse-hierarchy reformatting as a hallmark of experience-driven working memory plasticity.

## 1 Introduction

Consider an expert birdwatcher who can discern fine differences in the flight direction of a bird. As the glimpse is brief, a mental workspace needs to keep information online in the absence of sensory input so that it can be accessed at a later moment (Kiyonaga & Serences, 2025; Sreenivasan & D’Esposito, 2019). This ability, known as visual working memory, has been classically investigated using delayed match-to-sample tasks in macaques and humans, in which a stimulus is maintained across a delay and compared with a subsequent probe (Fuster & Alexander, 1971; Harrison & Tong, 2009).

A delayed comparison task places two demands on the visual system. First, the remembered direction must coexist with incoming input. Second, it must be maintained with sufficient precision to support fine discrimination. The contents of visual working memory can be decoded from sensory cortex during the delay period, independent of the level of BOLD activity (Riggall & Postle, 2012; Serences et al., 2009). In early sensory cortex, sensory and mnemonic information coexist in separable coding subspaces (Libby & Buschman, 2021; Rademaker et al., 2019). This arrangement, termed region-wide multiplexing, provides a “local comparison circuit” in which a maintained representation and a current percept can be read out and compared within shared circuitry (Hallenbeck et al., 2021; Rademaker et al., 2019). However, when mnemonic and sensory signals share the same circuit, incoming input can interfere with the maintained representation (Bettencourt & Xu, 2016; Rademaker et al., 2019). Rotation of the mnemonic representation into a subspace orthogonal to the sensory axis has been proposed as a mechanism that reduces interference (Buschman, 2021; Libby & Buschman, 2021; Xu, 2024).

Whether the memory retains sufficient precision for fine discrimination depends on the format in which it is coded and the level of the visual hierarchy at which it is maintained. Converging evidence suggests that memory representations become abstract during maintenance (Kwak & Curtis, 2022; Li & Curtis, 2023; Yu et al., 2020). For example, Chunharas et al. (2024) showed that visual memories become increasingly abstract across the visual hierarchy. Such abstract representations support stability and generalization but may be insufficient when precise stimulus information is required (Bettencourt & Xu, 2016; Duan & Curtis, 2024). Fine discrimination, however, depends on sufficiently precise representations of stimulus-specific information (Bays et al., 2024), and categorical representations can distort fine-grained differences in visual memory (Bae et al., 2015).

To address this question, we combined psychophysicspreserving stimulus-specific information, perceptual learning provides an ideal framework for examining how training reshapes mnemonic representations. The precision of fine discrimination improves with practice, a phenomenon known as perceptual learning (Fahle & Poggio, 2002; Watanabe & Sasaki, 2015). In delayed discrimination tasks, the maintained representation must preserve the information necessary to achieve these improved levels of precision. A central but unresolved question is how perceptual expertise reshapes the neural code that maintains visual information across delays: where in the cortical hierarchy and in what format training reorganizes mnemonic content. More specifically, a multiplex code can be characterized by the format of the maintained information, by the temporal coupling between sensory and mnemonic signals, and by the geometric relationship between their coding axes. Although perceptual learning can shift the contribution of cortical areas involved in delay-period maintenance (Jia et al., 2021), whether it changes the representational format of mnemonic codes remains unknown.

To address this question, we combined psychophysics with fMRI-based multivariate pattern analyses to characterize how perceptual learning reshapes mnemonic representations across the visual hierarchy. Participants underwent five days of intensive training on a fine motion-direction discrimination task, allowing us to track changes in delay-period codes before and after learning. We found that perceptual learning enhanced visual working memory performance by reformatting multiplex codes across cortical levels: delay-period representations in V1 became more sensory-like and closely aligned with sensory representations measured in an independent localizer, whereas mnemonic-format representations in V3A were reduced. Moreover, learning increased temporal coupling between sensory and mnemonic codes in V1 while preserving their near-orthogonal coding geometry. Together, these findings reveal that perceptual learning reorganizes working memory representations along a reverse visual hierarchy, shifting maintenance toward high-fidelity sensory-like codes that support fine delayed discrimination.

## 2 Results

### 2.1 Behavioral results

We trained participants over five consecutive days (∼3,000 trials in total) on a fine motion-direction discrimination task, in which two random dot kinematograms (RDKs) were presented for 0.2 s, separated by a 0.6 s interval (Fig. 1A-B). Across training sessions, discrimination thresholds gradually declined, amounting to a 35% improvement (Fig. 1C). Before and after learning, we assessed their thresholds in a motion discrimination task with a delay period of 2.4 s. Discrimination thresholds decreased at both the trained and untrained directions (both *t*(19) > 5.94, *p* < .001, paired *t*-tests). The reduction in thresholds was significantly larger at the trained direction than at the untrained direction (*t*(19) = 2.43, *p* = .025, paired *t*-test; Fig. 1D).

**Figure 1.**
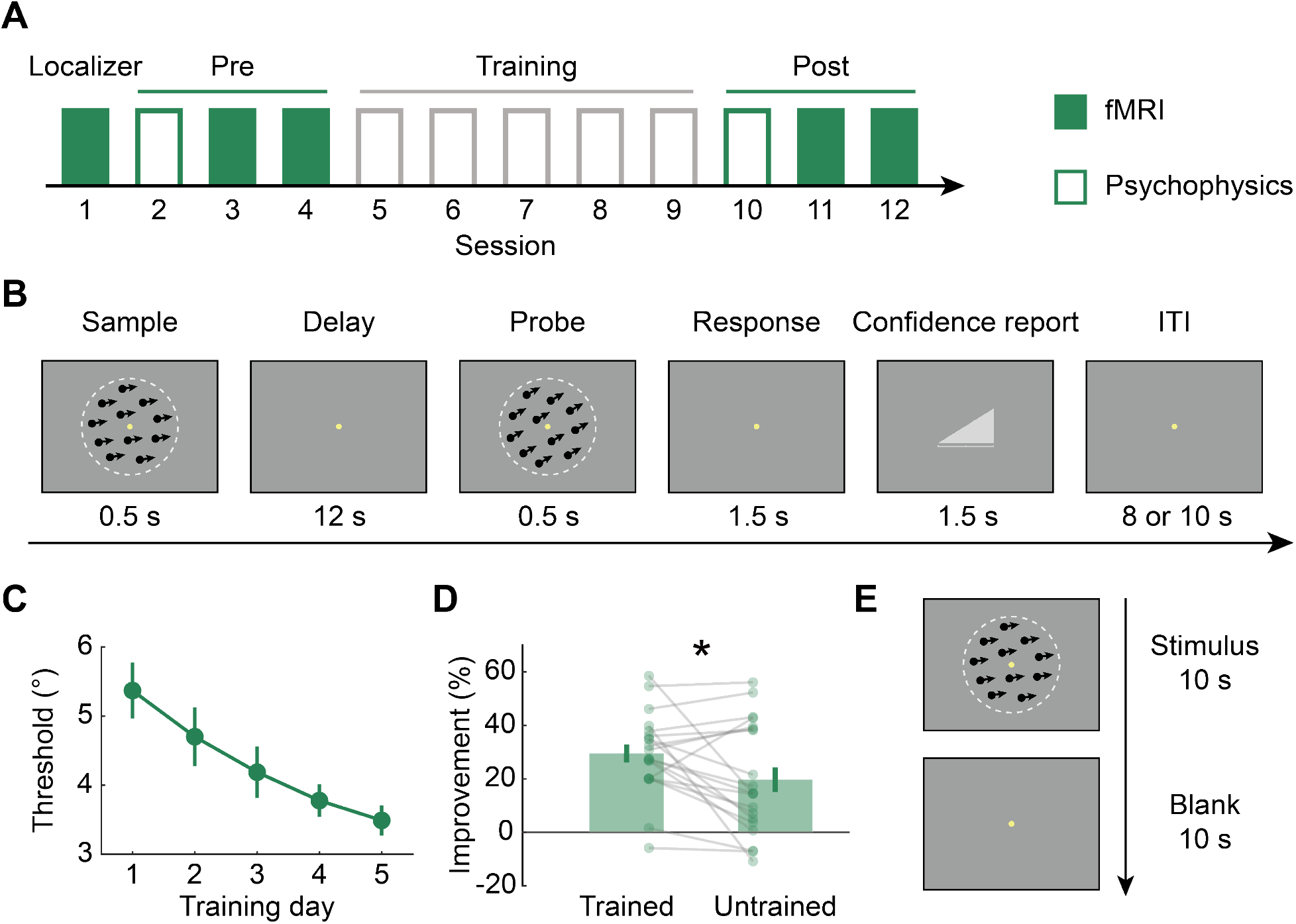
Protocol and behavioral results. (A) Experimental protocol. Subjects underwent five daily training sessions. The pre-training test (Pre) and post-training (Post) tests took place on the days before and after training. (B) Schematic illustration of the delayed motion discrimination task. Two random dot kinematograms (RDKs) were each presented for 0.2 s in psychophysical tasks, or 0.5 s during fMRI scanning, separated by a 2.4 s or 12 s delay. Participants judged whether the probe’s direction was clockwise or counter clockwise relative to the sample. (C) Learning curve. (D) Percentage improvement after training = (Threshold_Pre_ − Threshold_Post_)/Threshold_Pre_ × 100%. Error bars indicate ±1 SEM across subjects. *: *p* < .05, paired *t*-test. Dots and grey lines correspond to individual values. (E) Schematic illustration of the localizer. Stimulus blocks were each presented for 10 s, interleaved with 10-s blank intervals. Participants performed a fixation task.

### 2.2 Different coding formats of sensory and mnemonic representations

#### 2.2.1 Univariate results

Following each behavioral session at Pre and Post, participants performed the delayed motion direction discrimination task in fMRI scanner using a slow event-related design. Deconvolution on fMRI timeseries revealed a peak response at 5–6 s after sample onset (TR 3), followed by a decay period (7–14 s post-sample onset, TRs 4–7) before the rising response to the probe across ROIs (Fig. 2A). After training, a reduction in averaged response magnitude was found in V3A, both to the sample and during the delay (*p* < .05, permutation test) (Fig. 2A).

**Figure 2.**
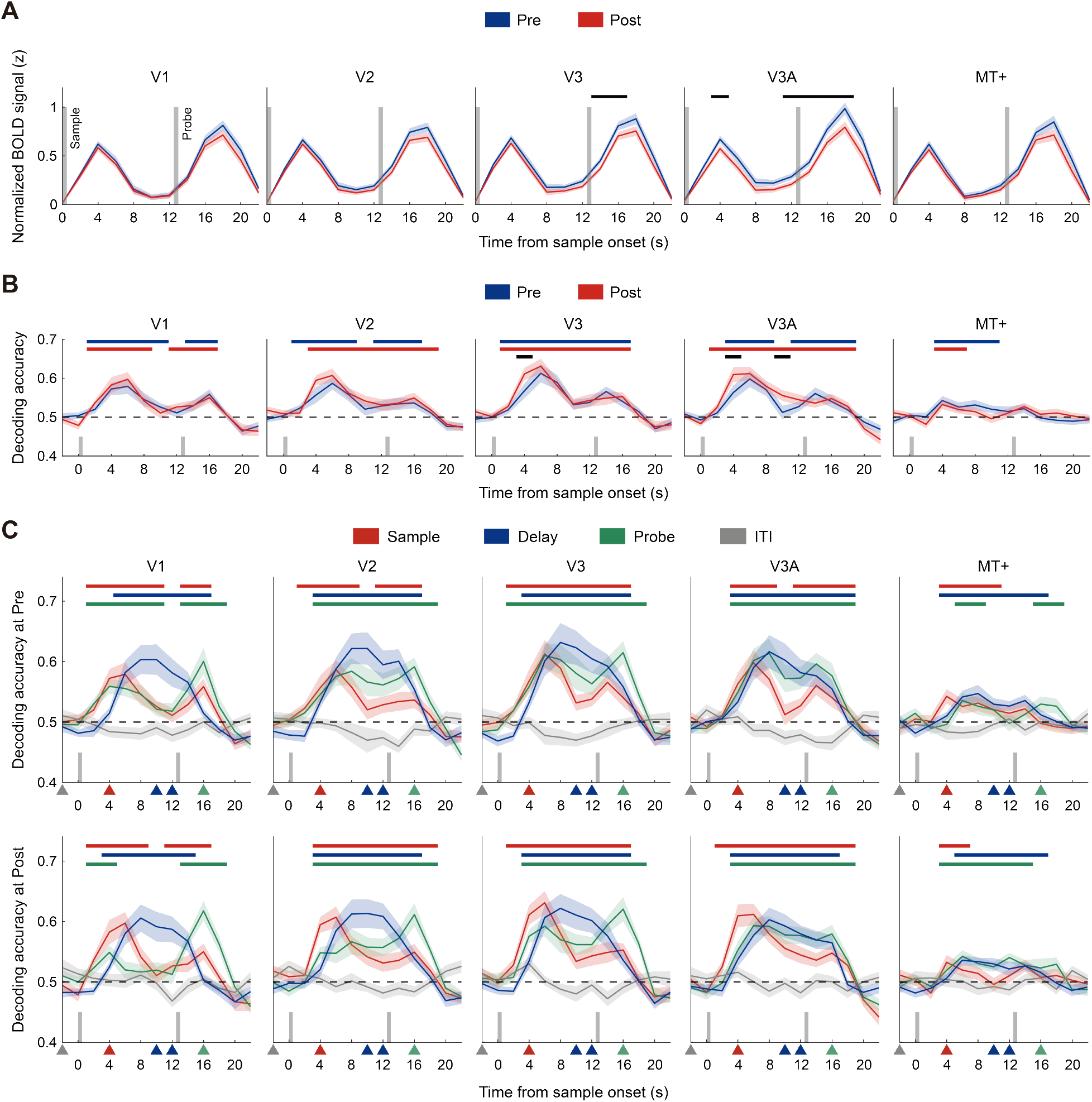
Univariate and multivariate results. (A) BOLD time courses in visual ROIs at Pre and Post. (B) Decoding time course from the “sample” classifier, trained on sample-evoked responses (TR 3) at Pre and Post. See TR-by-TR decoding time courses in Fig. S1. (C) Decoding time courses from four classifiers (Sample, Delay, Probe, ITI) for Pre (top row) and Post (bottom row) sessions. Colored triangles on the x-axis indicate time points used for training the corresponding classifier. Horizontal colored bars mark times when decoding accuracy exceeded chance (*p* < .05, cluster-based permutation tests). Black bars denote significant Pre vs. Post differences (*p* < .05, permutations tests). Vertical bars on the x-axis indicate the presentation of the 1^st^ (sample) and the 2^nd^ (probe) stimuli. Shaded bands show ±1 SEM across subjects. Data are unshifted in time.

#### 2.2.2 Pattern classification results

To examine whether the early, sensory-based representation remains stable throughout the trial, we trained the classifier on signals at the peak of the sample-evoked response, and then used it to construct a decoding time courses of direction representation (Fig. 2B). Across the ROIs, while the mean decoding accuracy deteriorates over the delay period, it was significantly above chance (50%, *p* < 0.05, permutation test) for most of time points. This overall pattern suggests that sustained BOLD activity carries stimulus-related information. After training, in addition to a higher decoding accuracy for the time point at the sample onset in V3 and V3A (both *p* < .05, permutation tests), V3A exhibited enhanced decoding accuracy for the delay period (*p* < .05, permutation test) (Fig. 2B).

In addition to the Sample classifier, we trained three classifiers to dissect the coding formats. The Delay classifier was trained on signals on time points after the peak of the sample-evoked response and before the rise of the probe-evoked response (i.e., TRs 4-7). The Probe classifier was trained from TR 9, corresponding to the peak of the probe-evoked response. The final classifier ITI was trained on the TR before the sample onset. We tested each classifier on all time points (Fig. 2C).

Overall, the time course constructed from the Delay classifier showed elevated accuracy in the delay period with a single peak, whereas time courses constructed from the Sample classifier and the Probe classifiers showed elevated accuracy at both onsets of the sample and the probe stimulus, since the two near-angle motion stimulus shared activation representation. These results demonstrate different coding formats of sensory and mnemonic representations.

### 2.3 Learning-induced changes in sensory and mnemonic information

At each timepoint, we applied the Sample and Delay classifiers to measure sensory and mnemonic information, respectively. To quantify learning effects for trained versus untrained directions, we extracted the signed distance, i.e., projection length, of trial-wise activation pattern to the classifier hyperplane (Fig. 3C).

**Figure 3.**
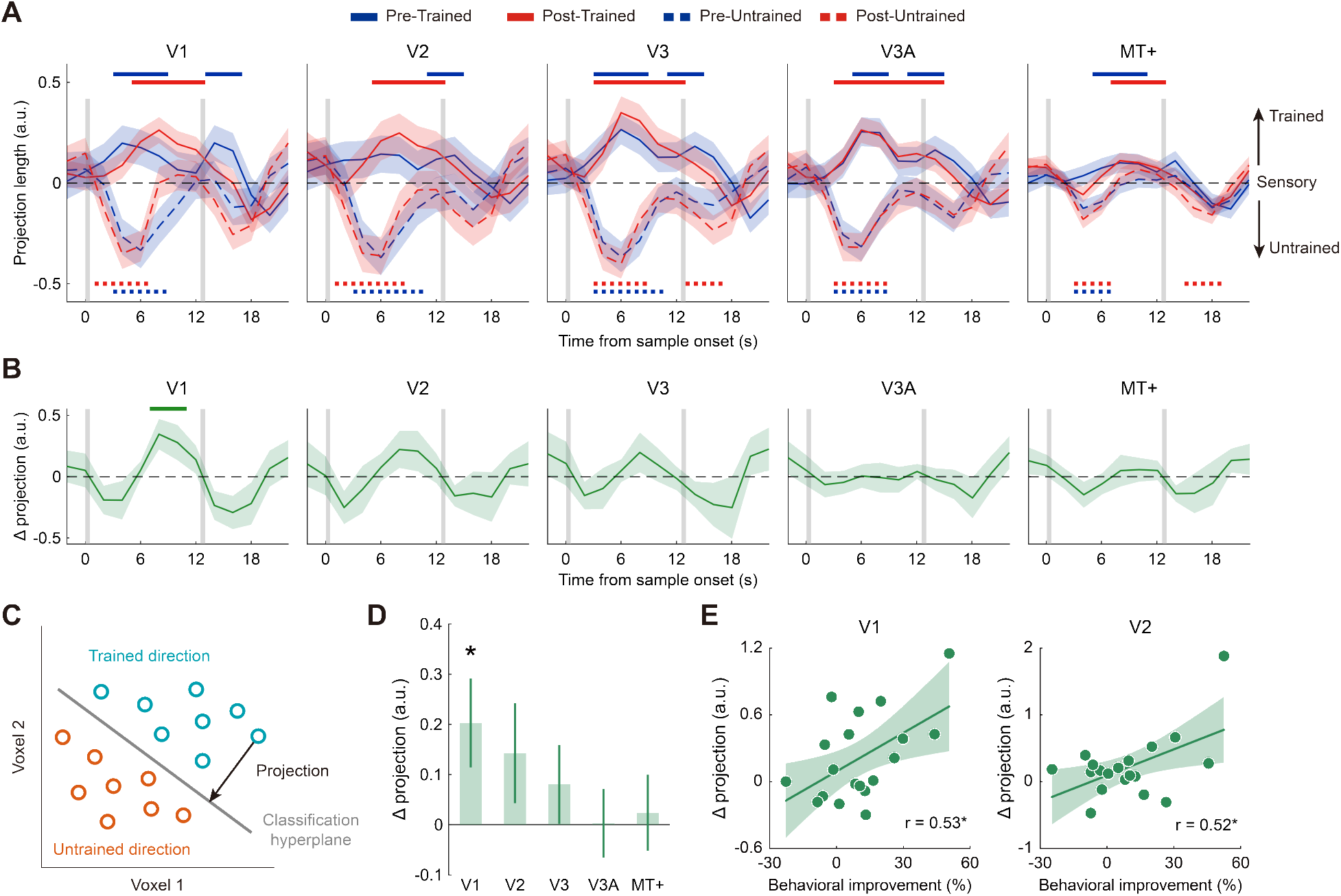
Perceptual learning enhances sensory information coding during maintenance in the early visual cortex. (A) Time courses of classifier-based sensory information. Positive values indicate classification strength towards the trained direction and negative values indicate classification strength towards the untrained direction. (B) Training-induced change in information. Shaded bands indicate ±1 SEM across subjects. Horizontal bars/dots at the top/bottom indicate the points at which the decoding information/change reached significance; *p* < .05, cluster-based permutation test. Vertical gray bars indicate the presentation of the 1^st^ (sample) and the 2^nd^ (probe) stimuli. (C) Schematic illustration of the linear classifier. The encoding axis of the classifier, i.e., the vector normal to the separating hyperplane, onto which voxel-wise activation patterns are projected to quantify the representational content of each ROI at each time point. Projections toward the trained direction were defined as positive. (D) Change in sensory information during delay period (TRs 4–7). Error bars indicate ±1 SEM across subjects. *: *p* < .05. (E) Correlation between the behavioral improvement and changes in sensory information in the early visual cortex. Solid lines denote best-fit linear regressions. Shaded bands denote 95% confidence interval.

For the time courses decoded from the Sample classifier (Fig. 3A), we examined the learning effect in sensory information by computing the Pre-Post difference and contrasted Trained versus Untrained direction at each timepoint (Fig. 3B). During the delay, sensory information significantly increased after training in V1 during the delay period (9-12 s post-target onset, *p* < .05, cluster-based permutation test).

Remarkably, the enhancement in sensory information in early visual areas predicted behavioral learning effect. Behavioral learning effect was quantified as the difference in percent threshold change between the trained and the untrained direction. Subjects with larger threshold gains exhibited larger enhancements in V1 sensory coding (r = 0.53, 95% CI = [0.12, 0.79], *p* = .015) and V2 (r = 0.52, 95% CI = [0.10, 0.78], *p* = .018; Fig. 3E).

Next, we examined learning-induced changes in mnemonic information in the time courses constructed from the Delay classifier (Fig. 4A). We observed a significant decrease in IPS0 at 9-16 s post-sample onset (*p* < .05, permutation test; Fig. 4B). Quantitatively, during the retention period (7–14 s post-sample onset), only V1 and IPS0 showed significant changes: sensory information increased in V1 (Δ = 0.21 ± 0.08; *p* < .05, permutation test), whereas mnemonic information in IPS0 decreased (Δ = –0.10 ± 0.05; *p* < .05, permutation test against zero). Interestingly, we identified a negative correlation between the change in sensory representation in V1 and the change in mnemonic representation in IPS across subjects (r = -0.47, *p* = .036, Spearman correlation; Fig. 4D). These results demonstrate that perceptual learning triggered dual changes in the delay period: an enhancement in sensory representation in V1 that goes hand in hand with a reduction in mnemonic coding in IPS.

**Figure 4.**
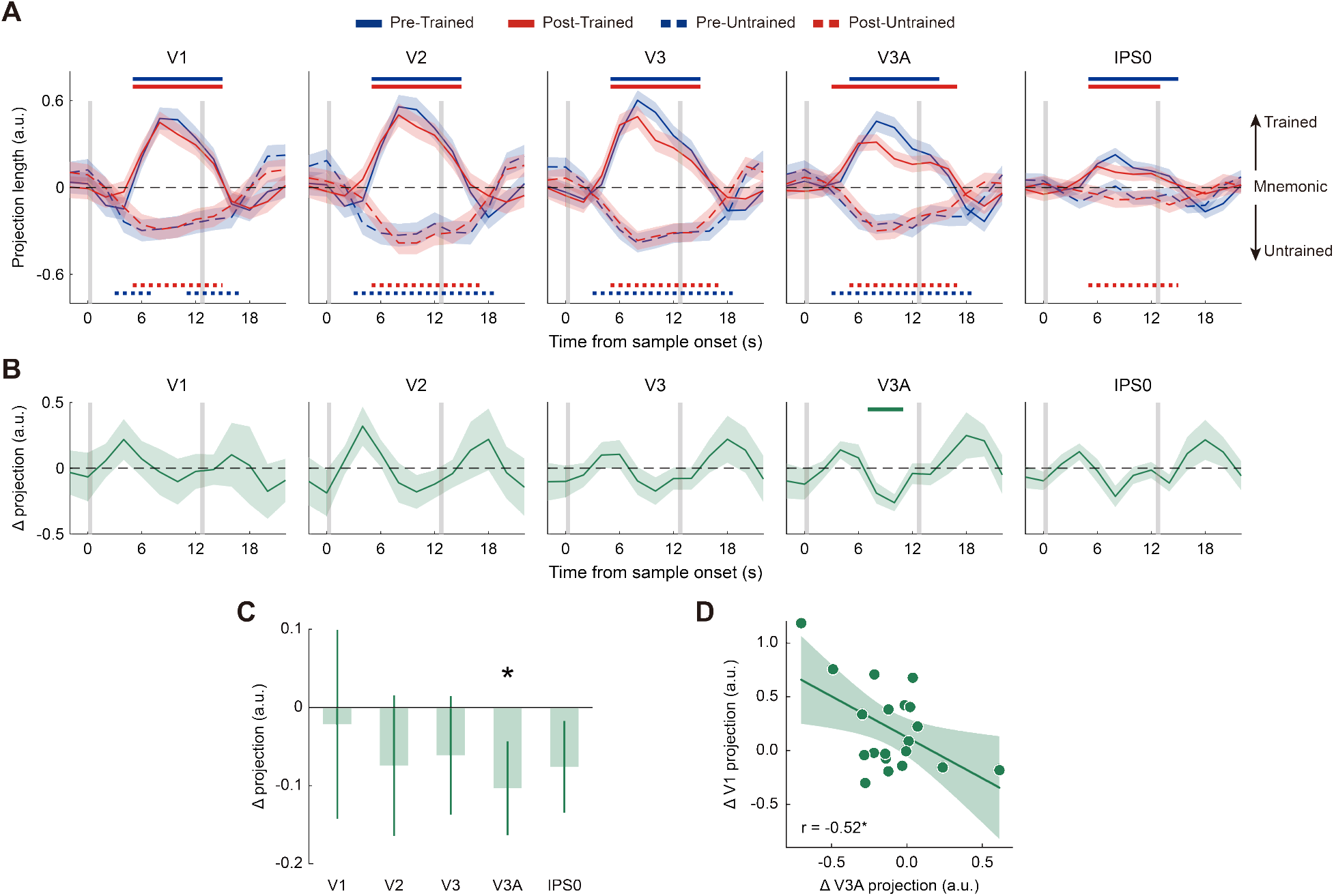
Perceptual learning reduces mnemonic coding in V3A. (A) Time courses of mnemonic information and (B) learning-induced changes in mnemonic information across visual and dorsal ROIs. Horizontal bars/dots at the top/bottom indicate the points at which the decoding information/change reached significance; *p* < .05, cluster-based permutation tests. Vertical bars on the x-axis indicate the presentation of the 1^st^ (sample) and the 2^nd^ (probe) stimuli. Shaded bands indicate ±1 SEM across subjects. (C) Change in mnemonic information during the delay period (4–7 TR post-sample onset) averaged across subjects. Error bars indicate ±1 SEM across subjects. *: *p* < .05. (D) Correlation between reduction of mnemonic information in V3A and increase of sensory information in V1. Dots represent individual subjects. Solid line denotes best-fit linear regressions. Shaded bands denote 95% confidence interval.

### 2.4 Learning promotes sensory-mnemonic alignment in V1

To further test whether the mnemonic codes become more sensory-like following training, we performed a representational similarity analysis to compare activation patterns between the main task and those from independent motion localizer runs (Fig. 1E). We computed Mahalanobis distances between BOLD patterns at each timepoint of the main task and each timepoint of the localizer, yielding a representational dissimilarity matrix (RDM). In the RDMs (Fig. 5A), the two salient diagonal “traces” (dark blue) correspond to the two sensory epochs triggered by the sample and probe stimuli. In between, there is a yellow region which corresponds to the delay epoch. The blue traces indicate that the stimulus-evoked pattern possessed high similarity across tasks, whereas the activation pattern during the blank interval differed, i.e., the activation during mnemonic maintenance period was different from the sensory trace in passive exposure condition.

**Figure 5.**
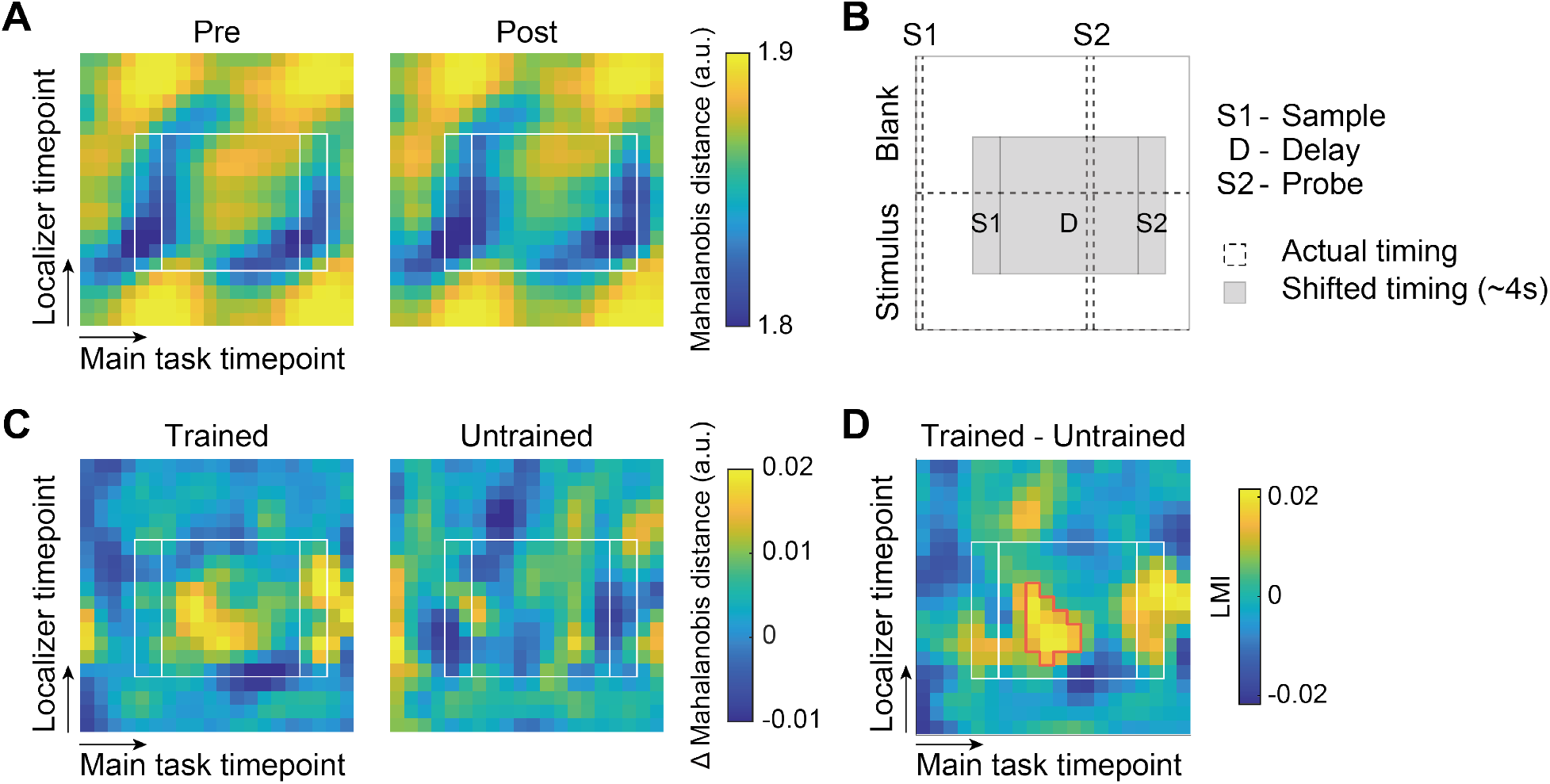
Perceptual learning aligns mnemonic and sensory representations in V1. (A) Averaged representational similarity matrix showing Mahalanobis distances between sensory localizer and main task in V1. Larger distance indicates greater dissimilarity. (B) Schematics of matrix plot. Dashed lines denote the actual timing of stimulus onsets and offsets. Grey boxes show the time windows shifted by ∼4 s hemodynamic lag. (C) Training-induced change (Pre − Post) in representational similarity for the trained (left) and untrained (right) motion directions in V1. Warmer colors indicate increased alignment between sensory and WM representations after training. (D) Learning modulation index (LMI) in V1. Red contours denote clusters exhibiting significant increases (*p* < .05, cluster-based permutation test). White dashed lines mark the onset and offset of each stimulus. White boxes denote time windows illustrated in (B). Data were smoothed to one-second resolution.

We computed the difference in dissimilarity matrices between Pre and Post for each condition (Fig. 5B). To isolate the learning effect specific to the trained condition, we further subtracted the distance change of the untrained direction from that of the trained direction, yielding a matrix of learning modulation index (LMI; Fig. S2). In V1, we observed a significant LMI spanning time points from 5 s to 12 s in the motion localizer and 7—14 s in the main task (red outline in Fig. 5C; *p* < .05, cluster-based permutation test). This enhanced alignment further demonstrates that mnemonic coding in V1 becomes more sensory-like after learning.

### 2.5 Alignment in sensory-mnemonic representational dynamics

Finally, we examined how sensory-mnemonic representational dynamics unfold over time by examining their joint evolution during the delay period. We embedded the moment-by-moment sensory and mnemonic decoding values into a two-dimensional information state space. Before training, the V1 trajectory initially moves rightward along the sensory axis after target onset while mnemonic information stays near baseline (Fig. 6A). It then curves upward-left as sensory information decays and mnemonic information rises, reflecting a transition from sensory to mnemonic code. At the onset of probe, sensory information drops to baseline but mnemonic information remains elevated. After learning, the trajectory shifts rightward across the entire interval, reflecting a sustained enhancement in sensory information; in contrast, mnemonic information during the delay period remains comparable to pre-training. Principal component analysis (PCA) demonstrates a change in the dominant joint dynamics following learning, as the first principal component (PC1) which captures dominant fluctuations in the joint sensory-mnemonic space differed significantly from Pre to Post (*p* < .05, permutation test; Fig. 6B).

**Figure 6.**
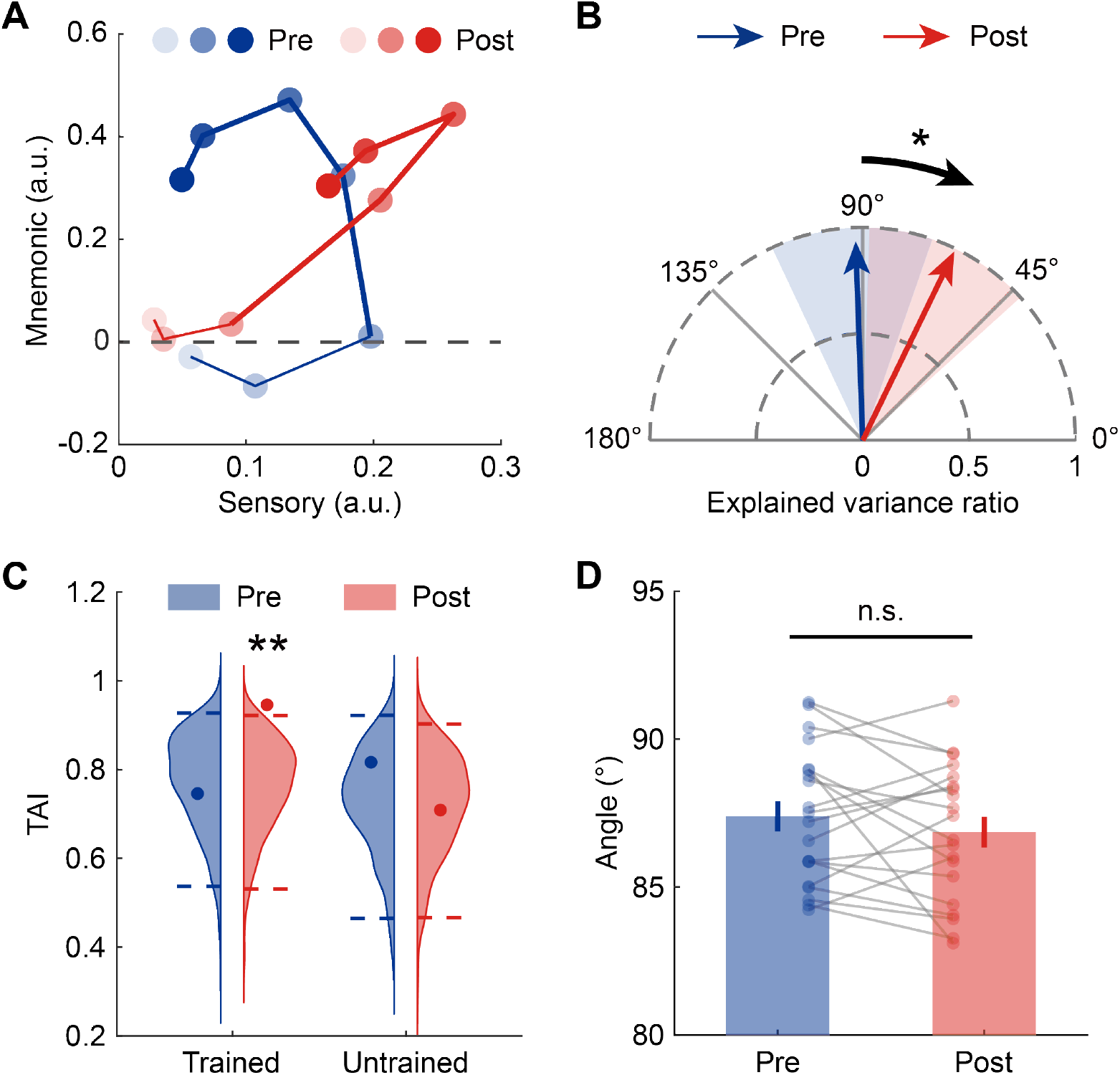
Dynamical alignment of sensory and mnemonic representations in V1. (A) Dynamic trajectories for trained motion direction at Pre and Post. Trajectories span from target onset to probe offset (0–13 s, 7 TRs in total). Each dot represents one TR. Dot color darkens over time; bold segments denote the delay interval. (B) First principal components (PC1) of dynamic trajectories in V1. Arrow directions indicate the PC axes; arrow lengths denote explained variance ratio. Shaded sectors denote the 95% confidence interval for direction of PC1 derived from 1,000 permutations. *: *p* < .05, permutation test. (C) Temporal alignment index (TAI) for trained and untrained directions in V1. A value close to 1 indicates a fully coordinated evolution of sensory and WM information. Violin plots show null distributions of index from 1,000 permutations; dashed lines mark the 95th percentile. Overlaid dots denote the empirical values. **: *p* < .01, permutation test. (D) Angle between sensory and WM axes of V1. Error bars indicate ±1 SEM across subjects. n.s.: not significant, paired *t*-test. Dots and grey lines correspond to individual values.

To evaluate the extent of joint evolution of sensory and mnemonic information across time, each momentary segment was projected onto the *y* = *x* direction to yield a vector component that quantifies the degree of alignment in temporal dynamics. For the trained directions, we identified a significant co-evolution at Post (*p* < .01, permutation test; Fig. 6C). These results suggest that learning synchronizes the transformation of sensory and mnemonic information.

In addition, we examined whether training altered the geometric relationship between the sensory and the mnemonic encoding axes (that is, the normal vectors of the classification hyperplanes). The angle remained near 90° in both sessions (Pre: 87.4° ± 0.51°; Post: 86.9° ± 0.52°) with no significant difference (*p* = .26, paired *t*-test, BF_10_ = 0.42; Fig. 6D).

## 3 Discussion

Using fMRI-based multivariate pattern analyses before and after intensive perceptual learning on a fine motion-direction discrimination task, we found that perceptual expertise reshapes both the neural locus and coding format of visual working memory. Learning enhanced high-fidelity sensory-format coding in V1, whereas the contribution of mnemonic-format coding in V3A decreased. The enhancement of visual sensory information in V1 correlated with behavioral learning improvement across individuals. Furthermore, sensory and mnemonic representations in V1 became more temporally coupled after training. These results indicate an experience-driven reallocation favoring the primary sensory cortex that is well suited for fine sensory comparison during delayed discrimination.

### 3.1 Experience-dependent reformatting across the visual hierarchy

First, we found that perceptual learning altered the distribution of maintenance-related codes within the visual hierarchy. Mnemonic-format coding decreased in V3A, a motion-sensitive area (Tootell et al., 1995) whose drection tuning could be sharpened by perceptual learning (Chen et al., 2015; Shibata et al., 2012). This reweighting emerged during mnemonic maintenance, with an increase in sensory-format coding in V1. The magnitude of these changes was negatively correlated across participants. These results indicate a reweighting of representational formats across visual areas, consistent with the reverse-hierarchy account, which proposes that perceptual learning progressively recruits lower-level sensory representations as task demands increase in precision (Ahissar & Hochstein, 2004; Sigman et al., 2005).

Converging evidence indicates that fine visual training can shift the contribution of cortical areas toward those best suited for the trained task. Learning can alter the relative contribution of cortical areas to perceptual decisions. In macaques, fine discrimination training changed the causal contribution of MT to depth perception (Chowdhury & DeAngelis, 2008), and in humans it shifted the cortical locus limiting performance from parietal to ventral visual cortex (Chang et al., 2014). The primary visual cortex provides a suitable substrate for fine motion judgments, as direction tuning is narrower in V1 than in MT (Albright, 1984; Lee & Lee, 2012; Snowden et al., 1992), and motion-direction training enhances early visual cortical responses, sharpens direction-selective representations, and strengthens feedforward interactions from early visual cortex to V3A (Song et al., 2024). The present results extend this pattern from perceptual processing to memory maintenance.

### 3.2 Plasticity of sensory-mnemonic multiplexing

A second question concerns how sensory and mnemonic codes are organized within V1 and how this multiplexing is modified by experience. Our data show that perceptual learning reshapes sensory–mnemonic multiplexing in this region. After training, V1 simultaneously carried stronger sensory-evoked responses during encoding and more sensory-like mnemonic signals during the delay. From a functional perspective, this coexistence may provide a substrate for a local comparison circuit, in which a maintained representation is compared with incoming sensory input (Rademaker et al., 2019). Alignment between mnemonic and incoming signals has been proposed to promote match detection through response gain (Serences, 2016).

This coexistence may be supported by the laminar architecture of early visual cortex, where feedforward inputs preferentially target middle layers, whereas superficial and deep layers participate in output and feedback pathways, respectively (Felleman & Van Essen, 1991; Nassi & Callaway, 2009). Future high-resolution fMRI studies could clarify how this mesoscopic organization of partially distinct laminar compartments supports sensory matching when mnemonic and incoming signals are aligned (Lawrence et al., 2018; van Kerkoerle et al., 2017; Voitov & Mrsic-Flogel, 2022).

### 3.3 Population coding dynamics and representational geometry

For naïve participants, sensory and mnemonic signals in V1 followed distinct temporal trajectories, with sensory information peaking around encoding and mnemonic information emerging later during the delay. This pattern is consistent with recent work showing that sensory and mnemonic representations can coexist within sensory cortex while remaining partially separable (Rademaker et al., 2019). Distraction can transiently reduce the fidelity of working memory representations in visual cortex, and trial-by-trial decoding biases in early visual cortex predict subsequent memory errors (Hallenbeck et al., 2021; see also Sprague et al., 2014). These findings indicate that working memory representations in visual cortex contribute to behavioral performance. Beyond coexistence, mnemonic representations may be reformatted to limit interference from concurrent signals. Under distraction, shifts in priority, or maintenance of multiple streams, memory codes can rotate away from sensory coding axes toward more orthogonal subspaces (Degutis et al., 2025; Libby & Buschman, 2021; Lorenc & Sreenivasan, 2021; Xu, 2024). Such separation may reduce interference between competing representations.

Our findings fit within this broader framework but reveal a different direction of reformatting. Perceptual learning did not shift the delay-period code in V1 away from sensory coding. Instead, training increased sensory-format information during the delay and promoted tighter temporal coordination between sensory and mnemonic signals. Thus, reformatting does not necessarily move memory representations away from sensory-like codes; its direction may depend on task demands. Working memory codes shift toward action-oriented formats when an upcoming response is specified (Bae & Chen, 2024; Henderson et al., 2022; Li & Curtis, 2023) and toward abstract formats when generalization is advantageous (Duan & Curtis, 2024; Kwak & Curtis, 2022).

Our results suggest that perceptual learning biases working memory reformatting toward the format most useful for the trained delayed discrimination. In our task, participants maintained a single motion direction for comparison with an upcoming probe. With one item in memory and no response prepared in advance, performance depended primarily on the precision of the maintained direction. This requirement favors a code that preserves small direction differences and matches the format of incoming sensory input. Consistent with this account, temporal coupling between sensory and mnemonic signals increased after training, while the angle between their encoding axes remained unchanged and close to orthogonal. This geometry permits coexistence with probe-evoked responses while preserving separability between sensory and mnemonic representations.

For an untrained observer, delayed discrimination may require comparison between a sensory representation and a transformed mnemonic representation. The sample is initially encoded in sensory cortex, and its representation may shift toward more transformed formats during maintenance (Chunharas et al., 2024; Kwak & Curtis, 2022). Our data suggest that training reduces this representational mismatch by strengthening sensory-format maintenance. After learning, sensory-format information in V1 remained elevated throughout the delay and became more temporally coupled with mnemonic representations, while their coding axes remained near orthogonal. This combination may allow fine stimulus information to be preserved while maintaining separability between concurrent sensory and mnemonic signals (Rademaker et al., 2019; Serences, 2016).

Interestingly, sensory and mnemonic coding axes remained near orthogonal after training, despite the increase in sensory-format information during maintenance. This preserved geometry suggests that perceptual learning enhanced the fidelity and temporal coordination of sensory-like maintenance without eliminating representational separation. In mouse auditory cortex, repeated exposure to temporally associated sounds rotated memory representations away from the sensory axis and reduced overlap between current input and memory of the preceding sound (Libby & Buschman, 2021). Thus, experience can reshape representational geometry, but the direction of reformatting depends on the computational demands of the task. In our data, sensory and mnemonic axes were already close to orthogonal before training, and five days of discrimination training did not measurably alter this geometry. This view is consistent with recent work emphasizing that working memory is intrinsically dynamic while maintaining a stable coding scheme (Wolff et al., 2020; see also Murray et al., 2017; Stokes, 2015). Related studies have shown that visual cortical representations evolve dynamically over time (Li & Curtis, 2023), and that the separation between working memory and distractor representations changes across time (Degutis et al., 2025). Building on this framework, our results suggest that experience selectively alters the strength and temporal coordination of sensory and mnemonic representations while preserving their distinct representational geometry.

In sum, we found that five days of training altered the balance between mnemonic and sensory-format coding across the visual hierarchy. Mnemonic-format coding decreased in V3A, whereas sensory-format maintenance increased in V1 and became more temporally coordinated with mnemonic signals. This pattern extends the reverse-hierarchy framework from perceptual encoding to memory maintenance. Importantly, learning strengthened the task-relevant sensory format while preserving the representational geometry that allows sensory and mnemonic codes to coexist. These findings suggest that perceptual learning reshapes working memory codes through a reverse-hierarchy reformatting process, strengthening high-fidelity sensory representations while preserving the distinct geometry of multiplexed codes.

## 4 Methods

### 4.1 Participants

Twenty healthy volunteers (12 female, 8 male) between the ages of 19 and 25 years (s.d. = 1.7) participated in the experiment. All participants provided written informed consent, had normal or corrected-to-normal vision, and received monetary reimbursement. The study was conducted at Tsinghua University. All procedures and protocols were approved by the Human Subject Review Committee of the Department of Psychology of Tsinghua University.

### 4.2 Stimuli and apparatus

The visual stimuli were random dot kinematograms (RDKs; number of dots: 900; diameter: 0.12°; speed: 6°/s; coherence = 1; luminance = 0.1 cd/m^2^) configured in a donut-shaped circular aperture with a 0.75° and 7° inner and outer radius, respectively. Stimuli were generated using MATLAB 9.10 (The MathWorks, Natick, MA) with Psychtoolbox-3 (Brainard, 1997). The stimuli were presented against a dark background (5 cd/m^2^) on an ASUS VG27AL1A 22-in liquid crystal display (LCD) monitor (resolution: 1920×1080; refresh rate: 60 Hz). from a viewing distance of 80 cm with their heads stabilized by a chin rest. In the fMRI scan, the stimuli were presented on a screen (refresh rate: 60 Hz; spatial resolution: 1920 × 1080; size: 88 cm × 50 cm) placed at the head-end of the scanner and subjects viewed through a tilted mirror from 200 cm. Throughout the experiments, participants were asked to fixate on a yellow dot with a diameter of 0.3° at the center of the monitor.

### 4.3 Design

The main experiment consisted of three phases: pre-training test (Pre), motion discrimination training, post-training test (Post). Pre and Post took place on the days before and after training (Fig. 1B).

During the training phase, each participant underwent 5 daily training sessions to discriminate motion directions around a direction *θ* (22.5° or 112.5°). One was designated as the *trained direction*, and the other as the *untrained direction*, with 0° defined as the rightward direction. In each trial, two RDKs (*θ* ± Δ*θ*) were presented sequentially for 200 ms each, separated by a 600 ms blank interval. Participants made a two-alternative forced-choice (2AFC) judgment of whether the direction of the second RDK relative to the first one (clockwise or counterclockwise). Responses were made via a self-paced keypress. Informative feedback was provided after each response, indicated by a change in the color of the central fixation point. The next trial began 1 s after the feedback. From trial to trial, *θ* randomly jittered within ±5° and Δ*θ* was adjusted using a QUEST staircase procedure to estimate the participant’s discrimination threshold at 75% correct (Watson & Pelli, 1983). The temporal order of these two stimuli was randomized. Each daily session comprised 15 QUEST staircases of 40 trials each. The discrimination thresholds derived from all staircases were averaged, and plotted as a function of training day.

During the test phase, psychophysical and fMRI tests were conducted using a delayed motion discrimination task. The psychophysical test was similar to that used in the training phase except that the two RDKs were separated by a 2.4 s blank interval and no feedback was provided. Each participant completed 6 QUEST staircases for the trained and untrained directions in a counterbalanced order. Prior to the experiment, participants practiced 2 staircases (i.e., 80 trials) for each condition to get familiar with the experimental procedure.

After acquiring the psychophysical thresholds in pre- and post-tests, each participant underwent 16 fMRI runs in two consecutive daily sessions (8 runs per session). In each trial, the first RDK, i.e., the target to be held in working memory, was presented for 500 ms, followed by a blank interval of 12 s, and then the second RDK (the probe) for 500 ms. The target direction was either 22.5° or 112.5°, with a random jitter within ±3° to ensure consistency during the delay period across trials. Participants were required to make a 2AFC judgment of the direction of the probe relative to the target (clockwise or counterclockwise), followed by a confidence rating and an inter-trial interval of 8 s or 10 s. Responses and confidence ratings were made within 1.5 s each, with no feedback provided. Each run consisted of 16 trials. The angular differences between the target and the probe were either 0.5× or 1×JND (just noticeable difference, determined as the discrimination thresholds in preceding psychophysical session).

Subjects completed eight block-design motion runs in an additional scan as localizer. Each run contained 8 stimulus blocks of 10 s, each for one of eight motion directions 22.5°, 67.5°, 112.5°, 157.5°, 202.5°, 247.5°, 292.5°, and 337.5°. Stimulus blocks were interleaved with 10 s fixation blocks. The size, coherence, luminance, and speed of the RDK stimuli were identical to those in the main task. Participants performed a fixation-color-change detection task throughout the run.

### 4.4 fMRI data acquisition and preprocessing

All scans were performed using a Siemens Prisma 3T scanner with a 64-channel head coil. Blood oxygen level dependent (BOLD) signals were measured with an EPI (echo planar imaging) sequence (slices = 72, voxel size: 2×2×2 mm^3^, TE: 34 ms, TR: 2000 ms, FoV: 200 mm, matrix: 100×100, gap: 0, flip angle: 90°). We also acquired distortion mapping scans with forward and reversed phase-encoding direction through the session to measure field inhomogeneities. A high-resolution (1×1×1 mm^3^) T1-weighted anatomical image was also acquired using an MPRAGE (magnetization prepared gradient-echo) sequence in the first scanning session.

Cortical gray-white matter volumetric segmentation of the high-resolution anatomical image was performed using Freesurfer’s recon-all script (version 6.0, https://surfer.nmr.mgh.harvard.edu). This anatomical image processed for each subject was the alignment target for all functional runs. The functional volumes were preprocessed using FSL (version 5.0, http://www.fmrib.ox.ac.uk/fsl), including motion correction, high-pass temporal filtering (0.015 Hz cutoff), and removal of linear trends. BOLD timeseries were Z-scored separately within run for each voxel.

### 4.5 Identifying regions of interest

Retinotopic visual areas (V1, V2, V3, V3A, and V4) were defined by a standard phase-encoded method developed by Sereno et al. (1995) and Engel et al. (1997), in which subjects viewed a rotating wedge that created traveling waves of neural activity in visual cortex. Subjects also performed a run to localize area MT+ (Chen et al., 2015; Huk et al., 2002), which contained eight moving dot blocks of 12 s, interleaved with stationary dot blocks of 12 s. MT+ was defined as voxels that responded more strongly to the moving dot blocks than to the stationary dot blocks within or near the occipital continuation of the inferior temporal sulcus. IPS0—3, SPL were defined using the probability atlas (Wang et al., 2015).

### 4.6 Event-related analysis

We performed a deconvolution analysis to estimate the hemodynamic response function (HRF) associated with the sample-onset and the following delay event for each voxel. This was done using a finite impulse response (FIR) function model consisting of a column marking the onset of the target stimulus, and then a series of temporally shifted version of that initial regressor in subsequent columns to model the BOLD response at each subsequent timepoint (Dale, 1999; Rademaker et al., 2019). In each ROI, we identified the top 80% of responsive voxels according to the peak value (5-6 s after sample onset, TR 3; Fig. 2A) of the estimated HRF. Voxels met this criterion in both Pre and Post sessions were selected for subsequent analyses.

### 4.7 Pattern classification analysis

Based on activation pattern from the selected voxels in each ROI, we trained linear SVM classifiers (Statistics and Machine Learning Toolbox, MATLAB 9.10, MathWorks, Natick, MA) to decode motion direction. Following a repeated random subsampling cross-validation procedure, all trials were randomly divided into equally sized training and test sets, each containing 32 trials for each motion direction. Four classifiers were trained using data from specific time windows: (1) the sample classifier, trained on response 5–6 s post-target onset (TR 3), corresponding to the peak response evoked by the sample; (2) the delay classifier, trained on averaged response 7–14 s post-sample onset (TRs 4–7); (3) the probe classifier, trained on response 17–18 s post target onset (TR 9), corresponding to the peak response evoked by the probe and response; and (4) the ITI classifier, trained on the TR before sample onset. The four types of classifiers were then tested on all time points in the left-out trials, yielding a time course of decoding accuracy across the entire trial. The cross-validation procedure was repeated 1,000 times for both Pre and Post sessions.

To further examine the sensory and mnemonic information of the sample, the projection distances from the hyperplane were extracted and signed according to the training labels. Positive and negative values indicate the trained direction and the untrained (orthogonal) direction, respectively.

### 4.8 Statistical test

We conducted statistical tests using MATLAB 9.10. Bayesian analyses were performed using an open-source toolbox (https://github.com/klabhub/bayesFactor). Paired sample *t*-tests and Bayesian paired *t*-tests were performed for psychophysical results. Bayes Factor (BF_10_) provides a ratio of the likelihood of the observed data in favor of alternative hypothesis (H_1_) over null hypothesis (H_0_).

The decoding timeseries was assessed using a cluster-based permutation test to correct for multiple comparisons over time (Maris & Oostenveld, 2007). First, we performed a one-sample *t*-test against 0 for each time point. We then identified contiguous clusters of time bins with uncorrected *p* values of *p* < .05 and calculated the sum *t* value for each cluster. A null distribution was generated by randomly flipping the sign of each participant’s information values across 1,000 permutations, and recording the maximum cluster-level statistic from each permutation. Observed clusters were considered significant if their test statistic exceeded the 95^th^ percentile of the null distribution (*p* < .05).

For the average change in classifier information, statistical significance was tested using nonparametric permutation tests. A null distribution was generated by randomly shuffling the sign of the information change for 1,000 times, and significance was determined by the proportion of null values exceeding the observed change (*p* < .05).

### 4.9 Representational similarity analysis

We computed Mahalanobis distances between voxel activity patterns elicited during a sensory localizer task and those elicited during the main task on a TR-by-TR basis (Fig. S3). For each motion direction, Z-scored BOLD signals from the localizer and from one run of the main task (each comprising 8 trials) were combined. Principal component analysis (PCA) was applied to this joint dataset, and principal components (PCs) cumulatively accounting for more than 80% of the variance were retained for estimating the noise covariance matrix (*S*). Following PCA, we averaged the PCs across trials, yielding two mean activation patterns for localizer (*x*) and main task (*y*), respectively. We then computed the Mahalanobis distance as:

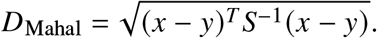

This procedure was repeated at each time point and for each run of the main task, and the resulting distances were averaged to yield a representational dissimilarity matrix for each individual. The RDMs were then averaged across subjects, and the resulting matrix was interpolated to a high temporal resolution (1,000 × 1,000; 20 ms resolution) using bicubic interpolation smoothing (Keys, 1981).

We computed the change in Mahalanobis distance between Pre and Post for the trained and the untrained direction:

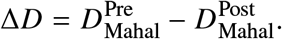

As Mahalanobis distance reflects representational dissimilarity, a positive Δ*D* indicates an increase in pattern similarity after training. To isolate the effect of perceptual learning, we defined the learning modulation index (LMI): LMI = Δ*D*_Trained_ − Δ*D*_Untrained_. Statistical significance of clusters within the LMI matrix was assessed using a cluster-based permutation test for each ROI.

### 4.10 Representational dynamics analysis

Sensory information and mnemonic information derived from the classifier-based projection method were assigned to X-axis and Y-axis, respectively, in a 2D state space. For Pre and Post, we averaged the time courses of sensory and mnemonic information across participants, resulting in seven (*x*, *y*) points corresponding to the TRs from sample onset (0 s) to probe onset (12.5 s). These time points formed a dynamic trajectory in the 2D state space.

To quantify the structure of each trajectory, we applied PCA to the seven 2D points from each session and ROI. The first principal component (PC1) captures the primary axis of variance in the trajectory, reflecting the dominant direction of representational change over time. The length of PC1 reflects the explained variance ratio.

To quantify the alignment of temporal dynamics between the two information. For each ROI, we first extracted the momentary vector during the delay. Each vector ***v***_*i*_ was defined as the difference between two adjacent TRs (bold segments in Fig. 5A). We then projected each vector onto the line of equal contribution from both information type (i.e., the diagonal *y* = *x*). For a vector ***v***_*i*_ = (*x*_*i*_, *y*_*i*_), the orthogonal projection onto the normalized diagonal ***u*** is given by

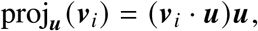

where

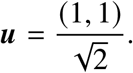

The length of the projected vector quantifies the degree of co-evolution between sensory and mnemonic information at that moment. The temporal alignment index (TAI) was defined as the mean magnitude of these projections across the delay period, ranging from 0 to 1. A value close to 1 indicates that sensory and memory information evolve in a fully coordinated manner along the diagonal axis, whereas values near 0 indicate that the dynamics are largely independent or dominated by one dimension.

Statistical significance of the learning effect was assessed using a permutation test. For each participant, Pre/Post labels were shuffled, preserving the temporal structure of each trajectory while removing consistent training effects. Null distributions were generated by repeating this procedure 1,000 times, averaging the pre-post difference across participants for each permutation.

### 4.11 Representational geometry analysis

To measure whether learning change the relative direction between sensory and WM encoding axes, we computed the cosine angle as:

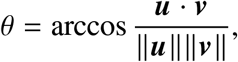

where ***u*** and ***v*** are the sensory and mnemonic information encoding vectors (that is, the normal vector of the hyperplane), respectively. We compared the angles between Pre and Post using a Bayesian paired *t*-test.

## Acknowledgements

This work was supported by MOST STI2030-2021ZD0203600, NSFC 32571227, and and Research Center for Brain Cognition and Human Development, Guangdong, China (Grant No. 2024B0303390003).

## Supplementary Materials

**Figure S1.**
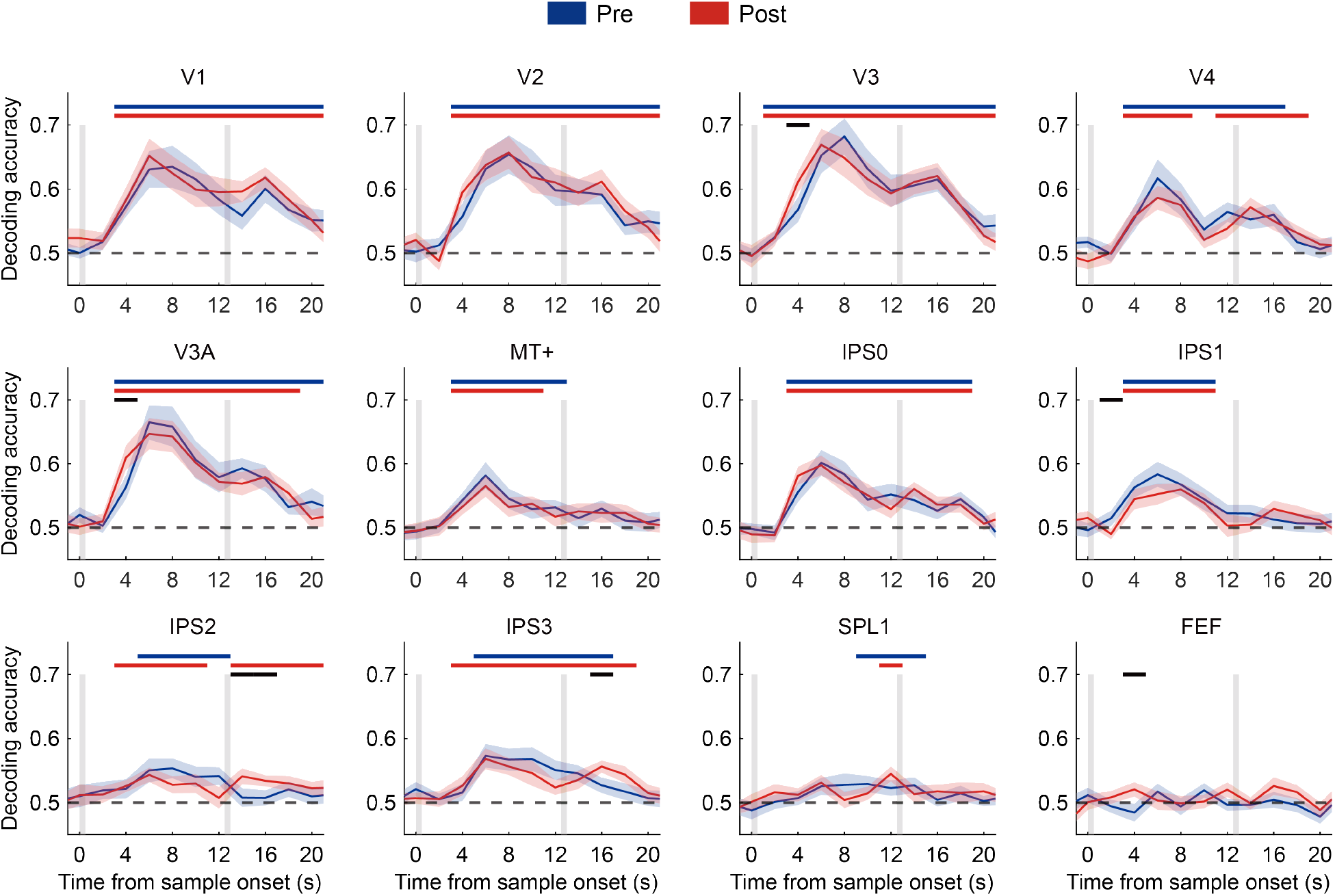
Decoding time courses for the main task of all ROIs. For each ROI, classifiers were trained and tested on BOLD signals from the same TR using a leave-one-run-out cross-validation procedure. This analysis was performed independently at each TR. Horizontal colored bars mark times when decoding accuracy exceeded chance (*p* < .05, cluster-based permutation tests). Black bars denote significant Pre vs. Post differences (*p* < .05, permutations tests). Vertical grey bars on the x-axis indicate the presentation of the 1^st^ (sample) and the 2^nd^ (probe) stimuli. Shaded bands show ±1 SEM across subjects. Data are unshifted in time.

**Figure S2.**
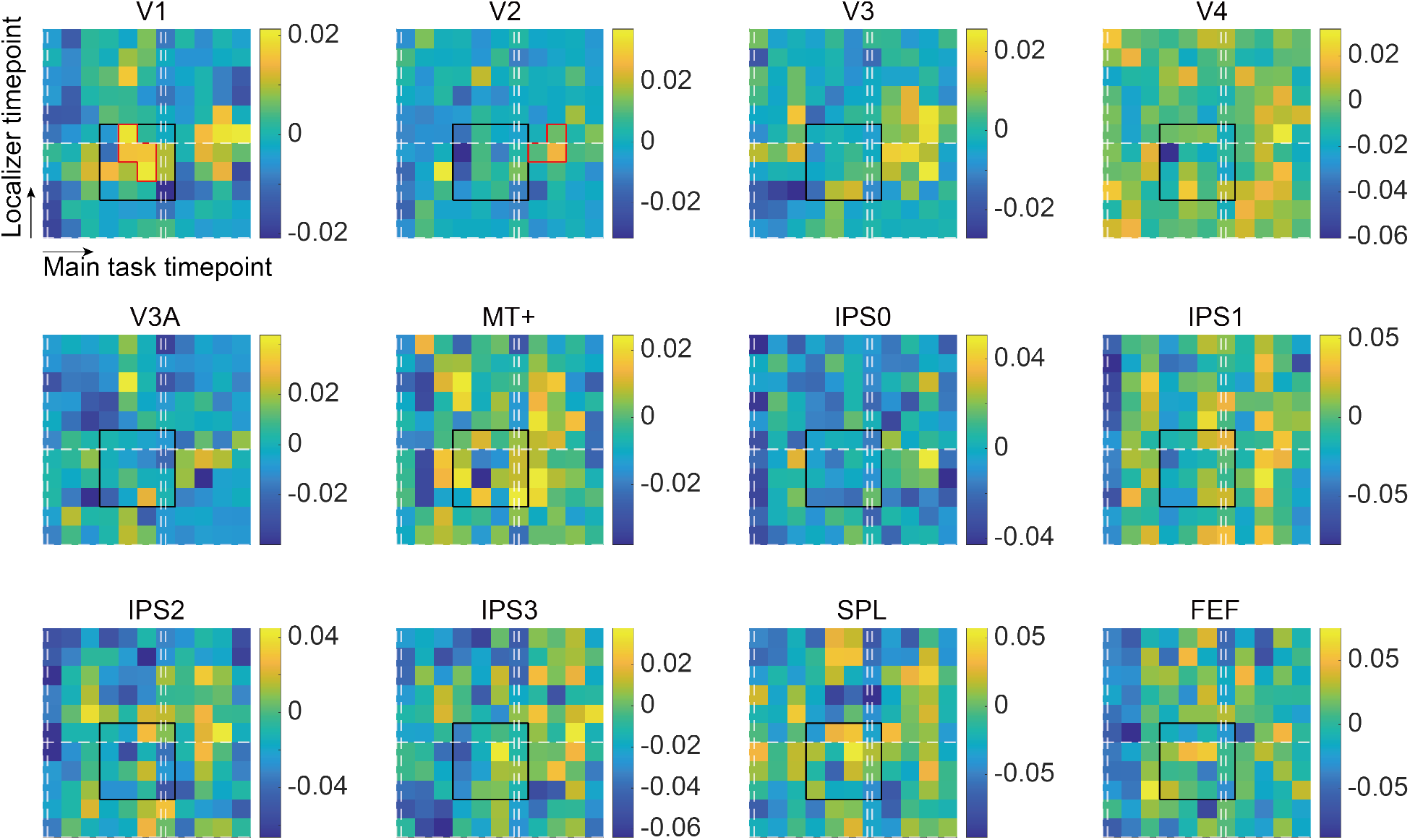
Learning modulation index (LMI) for all ROIs. (A) Red contours denote clusters exhibiting significant increases (*p* < .05, cluster-based permutation test). Solid black square highlights the temporal window spanning sensory encoding in the localizer (TRs 3-6) and WM maintenance in the main task (TRs 4-7). White dashed lines mark the onset and offset of each stimulus. Data were unsmoothed, i.e., each timepoint represents one TR (2 s). (B) LMI averaged over the defined time window in (A). Error bars indicate ±1 SEM across subjects. *: *p* < .05.

**Figure S3.**
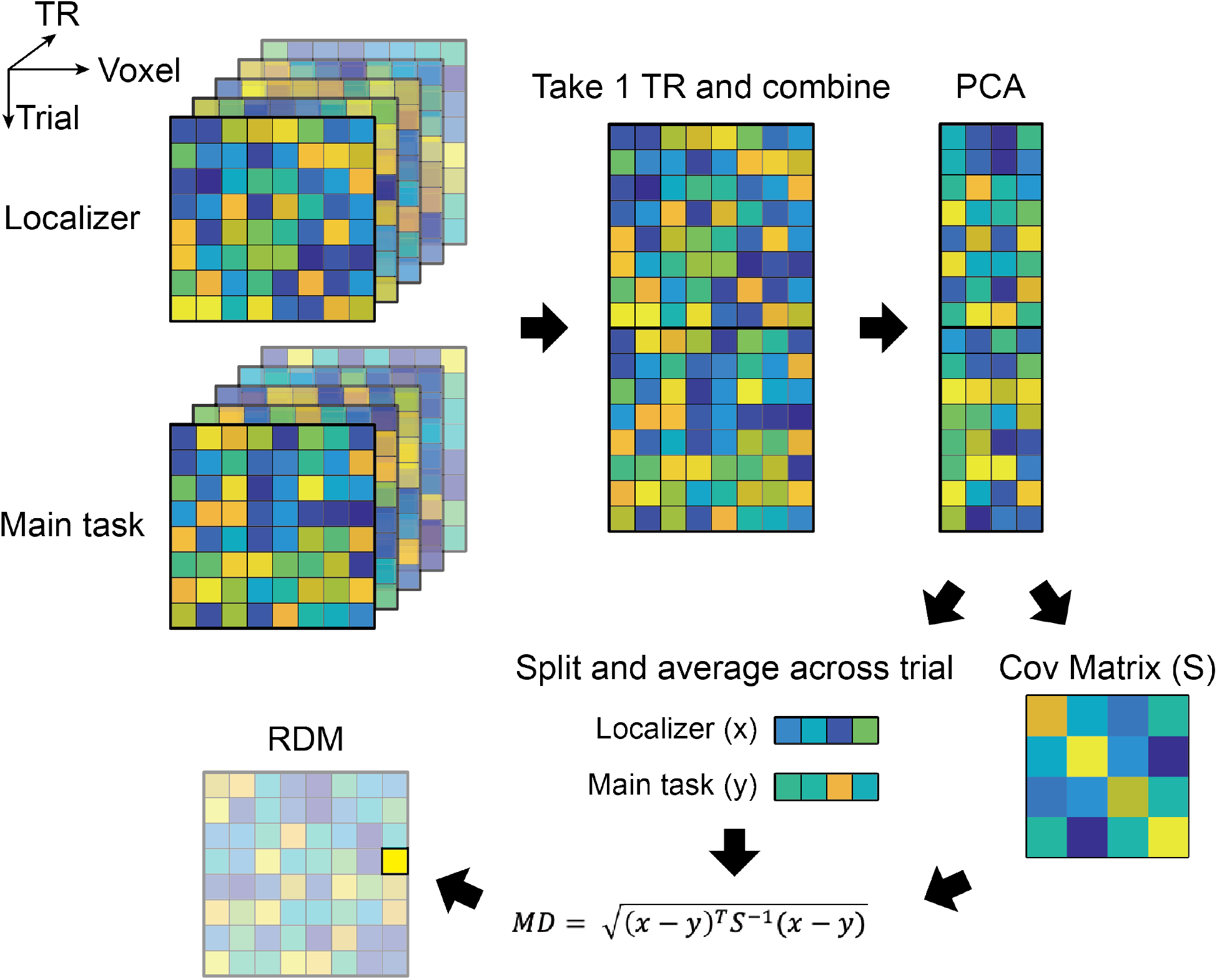
Schematics of computing representational dissimilarity matrix. We computed the Mahalanobis distance between the pattern induced by localizer and that in the main task on a TR-by-TR basis. For each motion direction, Z-scored BOLD signals from the localizer and from one run of the main task (each comprising 8 trials) were combined. Principal component analysis (PCA) was applied to this joint dataset, and principal components (PCs) cumulatively accounting for more than 80% of the variance were retained for estimating the noise covariance matrix (*S*). Following PCA, we averaged the PCs across trials, yielding two mean activation patterns for localizer (*x*) and main task (*y*), respectively. We then computed the Mahalanobis distance as: 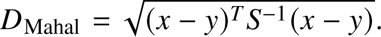 This procedure was repeated at each time point and for each run of the main task, and the resulting distances were averaged to yield a representational dissimilarity matrix for each individual.

## Notes

### Competing Interest Statement

The authors have declared no competing interest.

### Summary of Updates

Results updated; Fig. 1, Fig.4, Fig. 5 updated; Introduction, Discussion revised.

